# Analysis of Polygenic Score Usage and Performance in Diverse Human Populations

**DOI:** 10.1101/398396

**Authors:** LE Duncan, H Shen, B Gelaye, KJ Ressler, MW Feldman, RE Peterson, BW Domingue

## Abstract

Studies examining relationships between genotypic and phenotypic variation have historically been carried out on people of European ancestry. Efforts are underway to address this limitation, but until they succeed, the legacy of a Euro-centric bias will continue to hinder research, including the use of polygenic scores, which are individual-level metrics of genetic risk. Ongoing debate surrounds the generalizability of polygenic scores based on genome-wide association studies (GWAS) conducted in European ancestry samples, to non-European ancestry samples. We analyzed the first decade of polygenic scoring studies (2008-2017, inclusive), and found that 67% of studies included exclusively European ancestry participants and another 19% included only East Asian ancestry participants. Only 3.8% of studies were carried out on samples of African, Hispanic, or Indigenous peoples. We find that effect sizes for European ancestry-derived polygenic scores are only 36% as large in African ancestry samples, as in European ancestry samples (*t*=−10.056, *df*=22, *p*=5.5×10^−10^). Analyzing global populations, we show that relationships between height polygenic scores and height are highly dependent on methodological choices in polygenic score construction, highlighting the need for caution in interpreting population level differences in distributions of polygenic scores, as currently calculated. These findings bolster the rationale for large-scale GWAS in diverse human populations and highlight the need for better handling of linkage disequilibrium and variant frequencies when applying scores to non-European samples.

## Introduction

Awareness of the over-representation of participants of European ancestry in human genetics research has been broadly acknowledged^1–5^, and increasing the representation of diverse populations has recently become a higher priority for the research community^5–10^. This has led funding agencies such as the National Institutes of Mental Health to make genetic studies of diverse populations a priority. Accordingly, representation of non-European ancestry participants in genome-wide association studies (GWAS) increased, from 4% in 2009^1^ to 19% in 2016^3^. Most of the increase in non-European ancestry research is attributable to expansion of genetic studies of East Asian populations, as reported previously^3^ and as observed in our data (see below). As such, most populations are still severely underrepresented. This lack of representation, if not mitigated, will limit our understanding of etiological factors predisposing to disease risk, and will hinder efforts to develop precision medicine. It is also important to understand the implications of the European-centric bias of earlier genetic studies, for work that builds upon existing research. For example, researchers want to know how the limited diversity in early medical genetic studies impacts the use of polygenic risk scores in non-European ancestry populations.

The use of polygenic risk scores^11,12^ (PRS, also known as risk profile scoring, genetic scoring, and genetic risk scoring) has become widespread in biomedical and social science disciplines^13–15^. Businesses have commercialized this technology, including direct-to-consumer testing from 23&Me and other companies. Perhaps most importantly, there is hope that polygenic risk scores can improve health outcomes by accelerating diagnosis and matching patients to tailored treatments^16^. Polygenic scoring studies have demonstrated reliable, though modest, prediction using straightforward scoring methods^11,12^ and genetic data alone, for many complex genetic phenotypes (e.g. blood pressure^13,17^, height^18^, diabetes^9,19^, depression^7,20^, and schizophrenia^14^). Polygenic risk scores are calculated by summing risk alleles, which are weighted by effect sizes derived from GWAS results^11,12,21^. Commonly used methods account for ancestry using principal components (calculated on pruned genetic data). In the parlance of polygenic scoring studies, the *training* GWAS is referred to as the “discovery” sample, and the *testing* dataset is referred to as the “target” sample. No overlap between discovery and target datasets is imperative, as is the removal of related individuals from analyses, as demonstrated by Wray and colleagues^21^. Methods of prediction that offer modest improvements on this basic framework are also available^22–25^.

Polygenic scores can be constructed for any complex genetic phenotype for which appropriate GWAS (or other robust association) results are available. The challenges inherent in using polygenic scores – including modest predictive ability and considerations of statistical power in the interpretation of results – have been reviewed previously^21,26^. Recent research has focused on the generalizability of polygenic scores to non-European ancestry populations^27^. There is good reason to anticipate reduced predictive power in non-European samples because of differences in variant frequencies and linkage disequilibrium patterns between populations^12,28^. However, the expected magnitude of decreased performance is largely unknown. Few systematic studies of polygenic score performance across different ancestry groups are available, though see Hoffman and colleagues^13^ for a thorough investigation of blood pressure metrics. To date, all available Polygenic Scores in Diverse Human Populations information has been based on individual phenotypes or small numbers of observations^28,32^. Further, previous findings may need to be re-evaluated in light of newer findings about relationships between ancestry and GWAS results^29–31^. Thus, there is a need for systematic evaluation of polygenic score performance across multiple populations and phenotypes.

In addition to questions about the predictive performance of polygenic scores, a second major area of inquiry concerns differences in the distributions of polygenic scores for global populations, as currently calculated^27,30,31,33–39^. Multiple potential causes of observed distribution differences of polygenic scores have been reported, including drift^27^, selection^33,36–39^, artifactual differences due to uncorrected population stratification^30,31^ and different environmental effects^40,41^. We investigate relationships between global principal components and polygenic scores, and assess relationships between polygenic scores and phenotypes for height, using 1000Genomes^42^ data and country-level height information. These analyses will be helpful in calibrating researcher expectations about polygenic scores, as currently calculated, for diverse non-European ancestry populations.

## Materials and Methods

This study has two major parts. First, we analyzed data extracted from previous polygenic scoring studies in order to describe trends in polygenic scoring research and to provide the most comprehensive analysis of polygenic scoring performance available to date. Second, we analyzed properties of polygenic scores as calculated for the 1000Genomes individuals, and compared polygenic scores to country-level information about height. Relevant procedures for these two parts are described below. This work received a notice of determination that this was not human subjects research from Stanford University.

### Part 1: Extracting and analyzing data from previous polygenic scoring studies

We first identified studies with data suitable for extraction, via PubMed on January 23^rd^, 2018 using the following search terms: (Genome-Wide Association Stud* OR GWAS OR Genome Wide Association Stud*) and (polygenic risk score OR genetic risk score OR polygenic risk scor* OR genetic risk scor* OR risk profile scor* OR “genomic profile”). We sought to identify all polygenic scoring studies, of any complex genetic phenotype, from the first decade of polygenic scoring research. This yielded 1,226 studies, 733 of which were polygenic scoring studies (see **Figure 1**). From these 733 studies we extracted data about the ancestry of participants and methods of constructing polygenic scores.

We next identified studies that contained valid comparisons of the performance of polygenic scores in European ancestry participants and at least one other ancestry. Specifically, matched analyses (from two or more ancestry groups, from any given publication) had to use the same genotyping chip for all samples, the same weights for variants, the same algorithm for constructing polygenic scores, and the same methods of measuring phenotypes across all participants. Data from 29 studies met inclusion criteria. From these studies we then extracted effect size metrics for each ancestry group and then normalized score performance for all ancestry groups to performance within the European ancestry participants, by dividing all effect sizes (within each study) by the effect size of the relevant European ancestry sample, within each study. We multiplied values by 100 so that performance for each non-European ancestry sample could be expressed as a percentage of European ancestry performance, which was standardized to 100%. For example, the first polygenic scoring study of schizophrenia^12^ found that polygenic scores explained only 0.4% of phenotypic variance in an African ancestry sample, whereas 3.2% of phenotypic variance was explained in a matched European ancestry sample. Consequently, the value of 12.5% (100*(0.004/.032)) is represented as one yellow point in **Figure 2**, among the African ancestry observations displayed on the right side of the **Figure 2**. In **Figure 2**, each point represents one within-study comparison between a non-European ancestry sample and the matched (within-study) European ancestry sample. European ancestry performance is standardized to 100% in all comparisons and is represented by the blue circle on the left side of the figure. By normalizing within-study effect sizes to European ancestry effect sizes, we were able to combine observations across phenotypes, and therefore to obtain general estimates of polygenic score performance across ancestry groups and complex genetic phenotypes. We only analyzed results from studies employing the “classical” polygenic scoring approach, which includes “pruning and thresholding”, and allele weights derived from an independent discovery GWAS^11,12,21,26^. For construction of the boxplots using R, we used the default settings except for specifying that the whisker ends should reflect minimum and maximum values (‘range=0’).

### Part 2: Polygenic score properties for 1000Genomes individuals

For part 2, we used publicly available data from 1000Genomes^42^. Individual-level genotype data for 2,557 individuals was downloaded from ftp-trace.ncbi.nih.gov/1000genomes/ftp/release/20130502/. Weights for constructing polygenic scores came from publicly-available sources of GWAS results for height, body mass index (BMI), and schizophrenia^14,18,46,48^. Data about average human height, for countries of origin for 1000Genomes populations, was downloaded from a pre-compiled table with male and female heights by country: https://en.wikipedia.org/wiki/List_of_average_human_height_worldwide.

Data preparation and analysis for 1000Genomes samples: The full 1000Genomes dataset was first filtered to include only bi-allelic single nucleotide polymorphisms (SNPs) with greater than 0.1% minor allele frequency. In order to calculate principal components across 1000Genomes genotypes, we used second generation PLINK^53^ to obtain variants in approximate linkage equilibrium, and we also removed the MHC region of chromosome 6 (25-35Mb) and the large inversion region on chromosome 8 (7−13Mb). We then calculated 20 PCs across all individuals.

In order to obtain weights for constructing polygenic scores, summary statistics files (i.e. GWAS results) were pruned to include only variants in approximate linkage equilibrium, using second generation PLINK^53,54^ and the following thresholds: ‐‐clump-kb 500, ‐‐clump-p1 1, ‐‐clump-p2 1, ‐‐clump-r2 0.2. 1000Genomes European ancestry data was used as the source of linkage disequilibrium information for pruning all summary statistic files given the primarily European ancestry of discovery GWAS datasets.

Second generation PLINK^53^ was used to construct polygenic scores for each phenotype for the 1000Genomes participants, using 13 thresholds for including pruned discovery GWAS variants in scores, as follows: p<5×10^−8^, p<1×10^−6^, p<1×10^−4^, p<1×10^−3^, p<1×10^−2^, p<.05, p<.1, p<.2, p<.3, p<.4, p<.5, p<.75, p</=1. The statistical package R^55^ was used for plotting and statistical analyses (t-tests, correlations). Height phenotype data was downloaded from a compiled table of average heights, for males and females, by countries. Heights for males and females were averaged. Certain populations were excluded from the analysis of correlations between polygenic risk scores for height and height phenotypes for three reasons: Four populations were excluded due to lack of height phenotype data: Puerto Rican in Puerto Rico (PUR), Bengali in Bangladesh (BEB), Punjabi in Lahore, Pakistan (PJL), Mende in Sierra Leone (MSL). Two populations were excluded due to the combination of highly mixed country ancestry (impacting validity of height phenotype) and admixture of the 1000Genomes population (impacting variability in the polygenic scores for height): African Ancestry in Southwest US (ASW), African Caribbean in Barbados (ACB). One population was excluded due to the absence of a single European country of origin (needed for height phenotype information used in this report): Utah residents with Northern and Western European ancestry (CEU). Details are given in **Supplementary_Table_3_1000Genomes_countries of origin.xlsx**

## Results

### Usage and performance of polygenic scores in diverse human populations

How well different ancestry groups have been represented in the first decade of polygenic scoring research (2008-2017, inclusive) is shown in **Figure 1A**, which presents cumulative distributions of studies for specific ancestry groups across time. The field has been dominated by European ancestry studies. Across the 733 studies examined (see **Methods** for inclusion criteria and **Supplementary Table 1** for a list of studies), 67% included exclusively European ancestry participants. There have also been 140 studies conducted in exclusively Asian populations (19%), most commonly in East Asian countries (e.g., China and Japan). Only 3.8% of the polygenic studies from the first decade of polygenic scoring research concerned populations of African, Latino/Hispanic, or Indigenous peoples combined^*^. These results are similar to those reported by Popejoy and Fullerton^3^, who noted that non-European ancestry representation in GWASs was almost exclusively in Asian populations, and East Asian populations in particular.

By comparing representation of particular ancestry groups with world population estimates for those groups (**Figure 1B**), it is possible to quantify the over‐ or under-representation of each major ancestry group. European ancestry representation was approximately 460% of what it would be if representation was proportional to world ancestry. In contrast, African ancestry (17%) and Latino samples (19%) were under-represented relative to world populations. East and South Asian samples are combined in this figure, but it should be noted that representation of East Asian samples is much higher than South Asian samples, which have been included in very few polygenic scoring studies to date. Middle Eastern and Oceanic populations have the lowest representation in polygenic scoring studies relative to world populations for these groups (10% and 0%, respectively).

**Figure 1.**
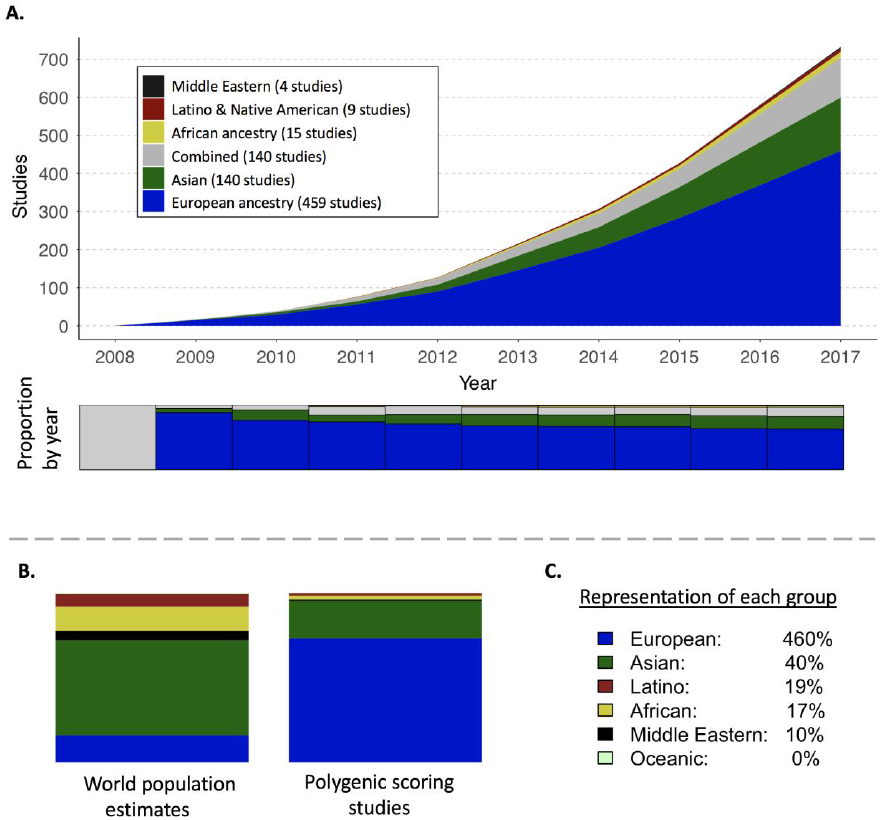
The first decade of polygenic scoring studies (2008-2017) focused primarily on European ancestry samples (N=733 studies). **A.** Cumulative numbers of studies by year are denoted by color. The stacked bar graph below the cumulative distribution plot shows proportional ancestry by year. **B.** Pie charts depict world ancestry representation (left) and polygenic scoring study representation (right). **C.** The percentage representation for each ancestry group is given, such that 100% would indicate equal representation in the world and in polygenic scoring studies. For example, European ancestry samples are over-represented (460%) whereas African ancestry samples are under-represented (17%).

Having analyzed the *use* of polygenic scores in different ancestry groups (above), we next assessed the *performance* of polygenic scores in multiple ancestry groups. Since most large-scale GWAS have been conducted in primarily (or exclusively) European ancestry individuals, our *a priori* hypothesis was that polygenic scores would perform best among European ancestry individuals, and less well for other populations. **Figure 2** provides an overview of polygenic score performance across ancestry groups. Results from all complex genetic phenotypes are analyzed together in order to increase the amount of data available for analysis.

Polygenic score performance was worst among African ancestry samples. The median effect size of polygenic scores in African ancestry samples was only 36% that of matched European ancestry samples (*t*=−10.056, *df*=22, *p*=5.5×10^−10^). Relative to matched European ancestry samples, performance was also lower in South (80%) and East Asian (93%) samples, but not significantly so. In sum, an expectation of poorer polygenic score performance in non-European ancestry populations seems reasonable given these data. Attenuation of predictive performances is likely to be most extreme in samples of African ancestry, consistent with, on average, greater genetic distance between European and African ancestry populations, than between European and other ancestry populations^28,45^.

**Figure 2.**
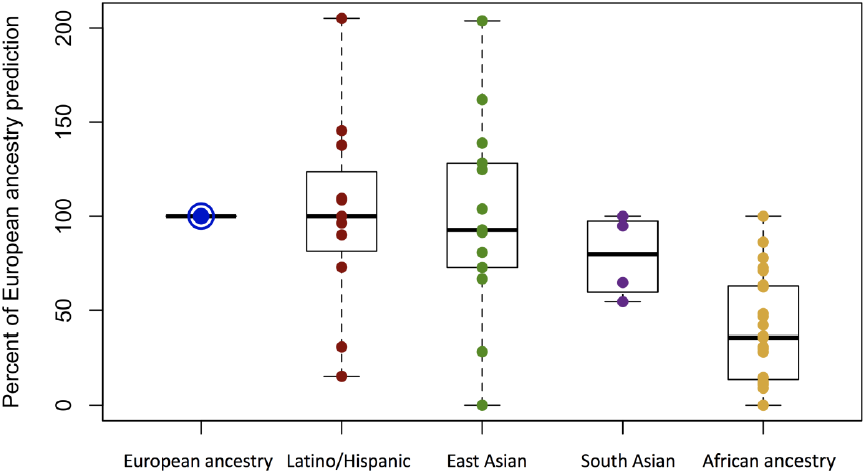
Performance of polygenic scores in Latino, East Asian, South Asian, and African ancestry samples, relative to performance in matched European ancestry samples (29 total studies). In order to make data comparable across studies, performance in each study was standardized to European ancestry performance, hence the single European ancestry y-axis value of 100%. Each point represents one pair of polygenic scoring analyses between a European ancestry sample and a matched sample from another ancestry (see text for details). Boxplots denote minimum, first quartile, median, third quartile, and maximum values.

### Correlations between global principal components (PCs) and polygenic scores, as currently calculated

We now consider questions about possible differences in polygenic scores among ancestral populations. Polygenic scores, as currently calculated, vary with ancestry. Indeed, polygenic scoring practices from as early as 2009 accounted for this^12^. The method used by Purcell and colleagues in 2009 (and frequently since) includes two steps for mixed ancestry samples. First, samples are separated into more ancestrally homogeneous subgroups (using visual inspection of plots of principal components calculated on all genetic data from all samples). Second, principal components are calculated again within each of these more ancestrally homogeneous subgroups, and are used as covariates in polygenic scoring analyses, which are conducted separately within each subgroup. **Figure 3** demonstrates why these ancestry analysis procedures have been used, given that many global PCs are strongly correlated with polygenic risk scores, as currently calculated.

**Figure 3.**
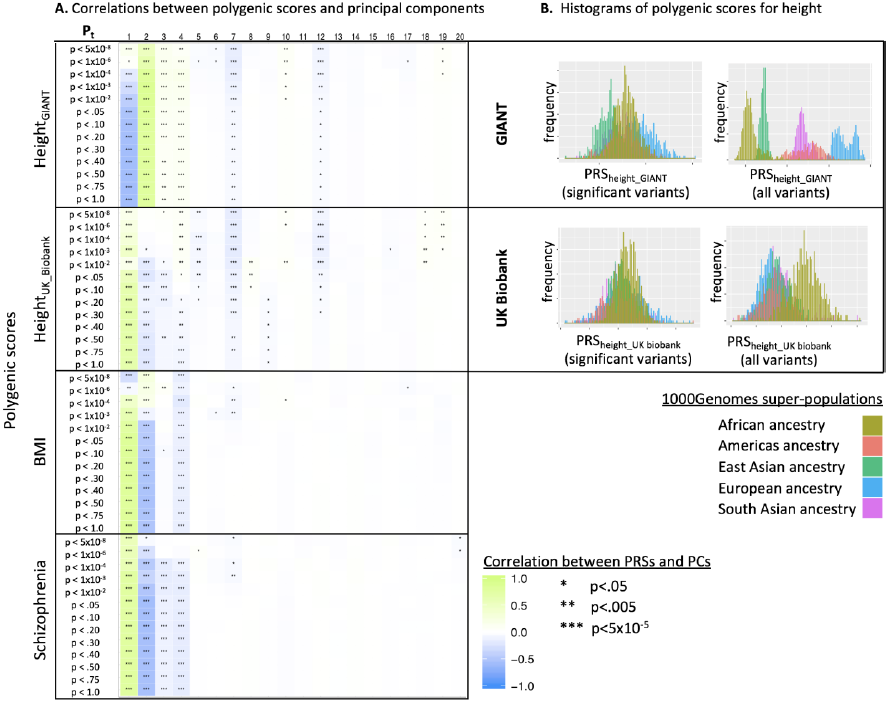
Properties of polygenic scores for 1000Genomes participants, across phenotypes (height _GIANT_, height_UK_Biobank_, body mass index, and schizophrenia) and across a range of p-values thresholds used to construct polygenic scores. **A.** Polygenic scores are often correlated with global principal components. For each phenotype, polygenic scores were constructed using 13 different p-value thresholds as applied to the discovery GWAS (denoted in the left-most column). Twenty global principal components (PCs) were calculated on 1000Genomes participants and are denoted across the top. Within the plot, correlations between each PC and each polygenic score are color-coded to reflect magnitude and direction of correlations from sky blue=−1 to lime=1. Stars indicate statistical significance of each correlation as follows: *p<0.05, **p<0.005, ***Bonferroni significant (p<5×10^−5^). **B.** Histograms of height polygenic scores for 1000Genomes participants are color-coded according to super-population. Two height GWAS were used to construct scores: GIANT (top) and UK biobank (bottom). Scores were constructed using two p-value thresholds: genome-wide significant variants (left) and using all variants (right). As can be seen for both GIANT and UK Biobank-based scores, the choice of p-value threshold applied to the discovery GWAS has a dramatic impact on score distributions among different populations. In general, inclusion of more variants in polygenic scores leads to greater differentiation among distributions of polygenic scores for global populations. P_t_=p-value threshold applied to discovery GWAS in order to construct polygenic scores, PRS=polygenic risk score, GIANT=Genetic Investigation of ANthropomorphic Traits, UK=United Kingdom, BMI=body mass index, PCs=principal components.

**Figure 3** shows that polygenic risk scores for the complex genetic phenotypes of height, body mass index (BMI), and schizophrenia are all significantly correlated with various global PCs (**3A**) and also that distributions of scores, as currently calculated, vary across global populations (**3B**). In Figure **3A**, it is primarily the first few PCs that are significantly correlated with polygenic scores, but non-consecutive and later PCs (e.g. 7 and 12 with height polygenic scores) may also be correlated. In order to construct Figure 3, we calculated polygenic scores and global PCs on all 1000Genomes individuals. We applied standard procedures including pruning and weighting alleles based on discovery GWAS results (see **Methods** for additional details). The results show that multiple, sometimes non-consecutive, PCs are strongly correlated with polygenic risk scores, as currently calculated. Many significant correlations are higher than *r*=.5, and 17.3% of correlations in **Figure 3** are significant after Bonferroni correction for 1,040 tests (i.e. 0.05/1,040= p<5×10^−5^; 20 PCs x 4 phenotypes x 13 p-value thresholds for each discovery GWAS = 1,040 tests). See **Supplementary Table 2** for correlations and corresponding p-values for **Figure 3A**.

In addition to showing significant correlations between global PCs and polygenic scores, as currently calculated, **Figure 3** also demonstrates that the choice of p-value threshold applied to the discovery GWAS (in the construction of polygenic scores) has a dramatic effect on score distributions across populations. The differences are so pronounced that the direction of the correlation between individuals’ values for a given PC and their polygenic scores may reverse across the range of p-value thresholds used to construct polygenic scores. For example, 1000Genomes participants’ scores for the first global principal component and their GIANT^18^-based polygenic scores for height are modestly positively correlated when only genome wide significant variants are used to construct scores (*r*=.14, *p*=3.4×10^−12^, faint green), whereas they are strongly negatively correlated when using all variants to construct polygenic scores (*r*=−.59, *p*=1.9×10^−223^, blue). In other words, there are many highly significant correlations, which vary not only in magnitude, but also in direction across the range of p-values used in the construction of polygenic scores. The effects of the methodological choice of p-value threshold on polygenic scores is further demonstrated on the right-hand side of **Figure 3B**, which shows the distributions of GIANT-based height polygenic scores for 1000Genomes participants, using two choices of p-value thresholds (left: genome-wide significant variants, right: all variants). Plots of distributions of UK Biobank-based^46^ polygenic scores for height are also shown on the right (bottom two plots).

**Figure 3** demonstrates key points relevant to the use of polygenic scores in diverse human populations. First, polygenic scores are often correlated with global PCs, and the correlated PCs are not necessarily consecutive (e.g. global PCs 1-4, 7, and 12 are correlated with height polygenic scores). Second, methodological choices of p-value threshold and discovery GWAS can have dramatic effects on polygenic scores, such that the magnitude and even the direction of observed relationships (e.g. between polygenic scores and PCs) may change across the range of commonly used parameters (e.g. across the range of p-value thresholds used to construct scores). These findings highlight the importance of treating ancestry properly in all analyses involving polygenic risk scores, and suggest that a conservative approach that analyzes polygenic scores separately in each ancestry group may be warranted, at least until a better understanding of polygenic score differences among populations (and across different phenotypes) is achieved. As noted by Chen and colleagues, explicit modeling of ancestry may afford even greater predictive power with polygenic scores^47^.

### Assessing putative correlations between global phenotypes and polygenic scores

Finally, we turn to the most difficult question: what causes differences in polygenic scores, as currently calculated, among different populations (e.g. see Figure 3B)? Differences could be real or artifactual (i.e. due to bias in data and/or methods), and five categories of explanations are listed below.

1) True differences due to drift
2) True differences due to selection
3) True differences in genetic effects due to environmental differences (gene-environment interactions)
4) Bias due to uncorrected population stratification in discovery and/or training samples
5) Bias due to discovery/training population data and/or polygenic scoring methods. Specifically, linkage disequilibrium (LD) structure and variant frequency are captured imperfectly with current methods (including genotyping and imputation), and they vary across populations, and currently available data resources are unequally representative of diverse global populations.

Drift has been implicated as an explanation for population differences in polygenic scores among populations^27^, but others have reported that drift is insufficient to explain such differences^33^. Further, initial estimates of the strength of polygenic selection on height in European ancestry populations^33,37^ have recently been greatly reduced^30,31^, based on findings of uncorrected population stratification in summary statistics from the GIANT Consortium^30,31^. There is also disagreement about whether or not differences in average polygenic scores among populations might contribute to differences in phenotypic values among the same populations (which could also be due to environmental variation). Some have noted apparent positive correlations between average polygenic scores and phenotypes for BMI^34^, lupus^35^, and height as calculated using GIANT Consortium scores^33,36,37^ and one group argued that there is no such correlation for height based on GIANT Consortium scores^27^. As described below, we include more data than used previously to address questions about potential correlations between global height polygenic scores and height phenotypes.

Using 1000Genomes data (as described in the **Methods**), we provide evidence consistent with artifacts contributing to differences in polygenic scores among global populations. In **Figure 4** we plot average polygenic scores for height of 1000Genomes populations on the x-axis, using three sources of weights for constructing scores (PRS=polygenic risk score):

- **4A** (top row) GIANT Consortium^18^ based scores: PRSheight_GIANT
- **4B** (middle row) UKBiobank^46^ based scores from the NealeLab: PRSheight_UKBiobank
- **4C** (bottom row) East-Asian GWAS based scores^48^ from He et al: PRSheight_EastAsian

On the y-axis, we plot average height for countries of origin for 1000Genomes populations, when available (see **Methods** for details and exclusions).

**Figure 4.**
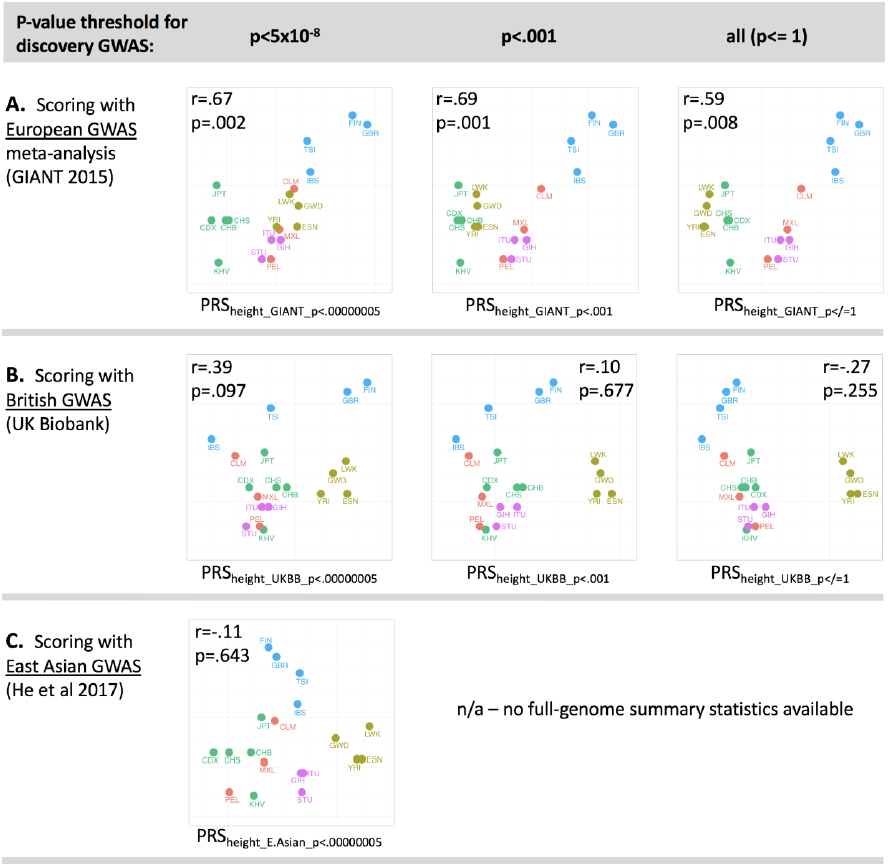
Scatterplots of height polygenic scores (x-axis) and phenotypic height (y-axis) show that correlations are not consistent across discovery GWAS. The y-values for phenotypic height are the same for each plot, and reflect average height of individuals in the country of origin or each population included. Three different GWAS of height were used to construct polygenic scores (i.e. three rows), and three p-value thresholds (i.e. three columns) were used for polygenic score construction applied to the relevant discovery GWAS. **A.** GIANT-based polygenic scores for height. **B.** UK Biobank-based scores for height. **C.** East-Asian-based polygenic scores for height. The last two plots are missing because only genome-wide significant variants were available for the East Asian GWAS of height^48^. GWAS=genome-wide association study, GIANT=Genetic Investigation of ANthropomorphic Traits, PRS=polygenic risk score, UK=United Kingdom, PCs=principal components. Population abbreviations are standard abbreviations from the 1000Genomes project^42,45^ and are included in Supplementary Table 3.

As shown in **Figure 4A**, height phenotypes for global populations (y-axes) are positively correlated with GIANT-based^18^ polygenic scores for height (x-axes), but not with UK-Biobank-based polygenic scores (**4B**) or East-Asian GWAS based polygenic scores (**4C**). Polygenic scores constructed using only genome-wide significant variants from GIANT (top left) were positively correlated with height phenotypes (*r*=.67, p=.002), as were scores constructed using larger numbers of GIANT-based variants (e.g. all variants, top right, *r*=.59, p=.008). Results in **4B** and **4C** demonstrate that correlations (or lack of correlations) between height and polygenic scores for height are dependent on discovery GWAS. Recent findings suggest that correction for population stratification may not have been adequate in GIANT^30,31^, and therefore the positive correlations observed in **4A** may be partially due to uncorrected population stratification. The dependence of correlation estimates on discovery GWAS is further illustrated in **4C**, in which the point estimate for correlation between height and East Asian GWAS based polygenic scores for height is negative (*r*=−.11, p=.643). Power in discovery GWAS is also relevant, and greater confidence should be attributed to the results in **4A** and **4B** because both European ancestry discovery GWAS were adequately powered to detect hundreds of height loci, whereas the East Asian height GWAS was only adequately powered to detect 17 loci.

These results suggest that both the ancestry of the participants in the discovery GWAS (e.g. European^18,46^ vs. East Asian^48^) and uncorrected population stratification^30,31^ contribute to observed positive correlations between GIANT-based polygenic scores for height and height across global populations. UK Biobank-based polygenic scores provide no evidence of a positive correlation between heights and mean height polygenic scores for populations. More research is needed to better understand the exact causes of differences in score distributions across populations and their putative relationships to phenotypes. Future research must also account for environmental effects on phenotypes, as well as variability in measurement validity and reliability across populations. Even for the relatively simple example of height (which is easily measured and for which major environmental influences are relatively well-understood) our analyses suggest that a great deal of caution should be used in drawing conclusions about polygenic score differences underling global phenotypic differences, until data resources are significantly improved (i.e. well-powered GWAS in diverse populations), and until a deeper understanding of relevant population genetics principles has emerged. As discussed further below, even more caution will be required for other phenotypes such as psychiatric disorders.

## Discussion

As conversations about personalized medicine make their way to the general public, it is important to recognize the need to include underrepresented populations in genetic studies. Among other concerns, the inclusion of participants representing diverse ancestries in research is imperative to ensure equitable benefit from scientific discoveries for diverse populations, and to prevent further increase in health disparities. Relevant to these longer-term objectives, our findings provide foundational information about polygenic risk score usage among diverse populations, summarized in four key points. First, polygenic scoring studies have primarily been conducted in European and East Asian ancestry populations. Second, the performance of polygenic scores in non-European populations is generally poorer than performance in European ancestry samples, and much worse in African ancestry samples. Third, polygenic scores for complex genetic phenotypes are often correlated with global principal components. Fourth, appropriate data resources are lacking to address most questions about putative differences in polygenic scores across global (non-European ancestry) populations. The straightforward, albeit expensive and time-consuming, solution to improving polygenic score performance across diverse populations is to create well-powered GWAS data resources for many different global populations.

Our finding that the predictive power of polygenic risk scores in poorer in non-European populations, particularly among African ancestry individuals, is almost certainly due to the considerably greater genetic diversity within many African populations, the phenotypic consequences of which is not well-captured by European-ancestry GWAS. Indeed, the relative impoverishment of genetic diversity among European ancestry and other non-African populations (due to past population bottlenecks) is a natural limitation of European ancestry GWAS. In addition to the important goal of collection of more samples from more populations, there can also be improvements in the manner in which variant frequencies and linkage disequilibrium are handled in polygenic scoring studies. The results presented here provide benchmarks for the relative performance of polygenic scores in diverse populations, as compared to performance in populations of European ancestry. This is important, because it not only informs power calculations for future research, but also highlights relative differences in predictive utility across diverse populations, which must be considered in public health calculations regarding when and how to use polygenic risk scores. Available data, confined mostly to European ancestry samples, provides a best-case-scenario upon which policy decisions may be based, rather than an accurate portrayal of how polygenic scores will work in ancestrally diverse patient samples.

Concerning expectations about polygenic score performance in non-European ancestry samples we note that polygenic scores for many complex genetic phenotypes are strongly correlated with global PCs, which highlights the critical importance of appropriate statistical methods for the analysis of genetic data from non-European ancestry populations. Testing future polygenic scoring results for robustness to the inclusion of variable combinations of PCs will reduce the chances for spurious results (that are actually attributable to ancestry). In sum, the preponderance of genetic studies based on European ancestry samples has led to a situation in which polygenic scores are approximately one-third as informative for African ancestry individuals, as they are for European ancestry individuals. This is presumably true for commercially available tests as well, and consumers should be aware of the differential performance of tests across individuals.

Regarding scientific and public perception of polygenic scores, it is important to address apparent differences in polygenic score distributions across populations. Our findings suggest that it is currently not possible to know precisely the distribution of polygenic scores for diverse non-European populations, for any complex genetic phenotype, because data resources for most populations are currently inadequate. Further, as we have shown, the ordering of population distributions of polygenic scores varies under accepted methods of constructing these scores (i.e. using different p-value thresholds for variant inclusion in scores and using alternative discovery GWAS). Explanations for these differences are currently incomplete. Until vastly superior data resources are available – including large scale GWAS in multiple globally representative populations – scientists are unlikely to reach consensus regarding the existence, nature, and exact causes of polygenic score differences among populations.

In our analysis of possible relationships between average phenotypes for global populations and average polygenic scores for those phenotypes in global populations, we chose to examine height because it is easily measured and because factors affecting height (e.g. nutrition) are also relatively easily quantified. In contrast, research on other variables such as weight, smoking status, psychological symptoms, and cognitive performance requires more careful control for environmental confounders (including variables like social status), which are often correlated with ancestry and therefore may also be correlated with global principal components and polygenic scores (as currently calculated). This means that confounding of environmental and genetic effects is likely. For example, social experiences such as being subjected to racism are prime candidates for confounding in genetic studies.

In closing, we emphasize the need to engage experts from other disciplines, such as social psychology and bioethics, as geneticists attempt to characterize genetic effects on complex genetic phenotypes impacted by social factors (especially psychological and cognitive variables). This is necessary because societal influences including socioeconomic status and discrimination can powerfully influence these phenotypes^52^, and these causal social factors often co-vary with ancestry. In genetic research, there is potential for relative blindness to non-genetic influences on phenotypes. Consequently, experts must be consulted in order to properly account for non-genetic influences on many complex genetic phenotypes and gene-environment interplay (i.e. correlations and interactions). Nevertheless, with cautious and broadly-informed research, potential medical benefits of correctly interpreting polygenic variation within and among populations can be realized.

## Acknowledgements

Pilot grant to LED and BD from the Stanford Center for Clinical and Translation Research and Education (UL1 TR001085, PI Greenberg) helped fund this work. The Stanford Center for Computational, Evolutionary, and Human Genetics, CEHG, supported this work.

## Author Contributions

LD and BD designed the project, LD and HS conducted the analyses, BG and KR contributed data, RP and MF provided critical feedback and framing, all authors contributed to writing of the manuscript.

## Materials and Correspondence

Correspondence and materials requests should be addressed to LD

## Footnotes

* Note that we retain populations names from the original reports (e.g. “Native American” and “Middle Eastern”) for Figure 1 in order to maintain consistency in terminology. “Combined” means that more than one ancestry group was included in the study (e.g. European ancestry and Asian ancestry participants).

